# A Non-Destructive Method for Quantifying Tissue Vascularity Using Quantitative Deep Learning Image Processing

**DOI:** 10.1101/2020.04.06.028555

**Authors:** Austin Veith, Aaron B. Baker

## Abstract

The quantitative analysis of blood vessel networks is an important component in the understanding and analysis of vascular disease, ischemia, cancer and many other disease states. However, many imaging techniques used for imaging vascular networks are time consuming, prevent further analyses or are technically challenging. Here, we describe a nondestructive technique for imaging the vessels in harvested tissue samples that relatively rapid and effective visualization of *in situ* vasculature. Furthermore, this technique allows for further analysis of the sample using histochemical staining, immunostaining and other techniques that can be performed on fixed tissue. The method uses ex vivo permeation of the tissue with an iodine solution and micro CT imaging to visualize the vascular network. Using a deep learning algorithm, we trained a convolutional neural network to automatically segment blood vessels from the surrounding tissue for analysis and create a map of the vasculature. While this method cannot achieve the resolution obtainable through destructive techniques, it is a simple and quick method for visualizing and quantifying vascular networks in three dimensions prior to analysis with conventional histological techniques. The global visualization of the three-dimensional vascular network provided by this method gives additional insights into complex changes in the vascular structure and can guide further histological analyses to specific regions of the vascular network. Overall, this method is a simple technique that can be added to most conventional tissue analyses to enhance the quantification and information derived from animal models with vascular network remodeling.

## Introduction

The quantitative analysis of blood vessel networks is a key aspect of many preclinical models of disease, including those for studying vascular disease^1^, ischemia^2^, angiogenesis^3^ and cancer^4,5^. While histological analyses allow the study of the tissues in detail, the overall complex blood vessel network structure is difficult and time consuming analyze using only two-dimensional sections of the tissue. One method for analyzing the vascular network is through the perfusion of the vascular network with a polymer that provide contrast and can be imaged at high resolution using X-ray Micro-CT Microtomography (Micro CT)^6-9^. This method provides three-dimensional vascular network with high resolution but also precludes further analysis of the tissues by histology or other methods. Another method has involved the use of tissue clearing techniques, immunostaining and light sheet microscopy to allow the 3D structure of tissues^10,11^. These methods can also provide images of the vascular network; however they are time consuming, require specialized equipment/imaging systems and also prevent many additional analyses from being performed on the tissues.

In this work, we have developed an optimized, non-destructive technique for tissue vasculature imaging, network segmentation, and quantification using deep learning methodologies. This method is relatively simple and can be added to the tissue analysis pipeline prior to performing histological analyses for virtually any animal model of vascular network remodeling. It provides an overall three-dimensional view of vascular network prior to histological analysis, adding insights into changes in the vascular network that may not be obvious from two dimensional tissues sections and affording a guide to targeted analyses to specific regions of tissue. A key aspect of the methodology is the use of a convolutional neural network^12-14^, which speeds the analysis the tissues to rapidly delineate vascular structures. Further, since it is nondestructive, it can provide additional insights without increasing number on animals used for the study. Overall, the method is a general means to quantify the vessels within the tissue in a relatively quick and non-destructive manner, while preserving the immune reactivity and tissue structure for further histological analysis.

## Methods

### Tissue harvest and processing

All experimental procedures were conducted according to animal care guidelines of the University of Texas at Austin Institutional Animal Care and Use Committee (Protocol ID AUP-2019-00328). Animal care was performed and humane endpoints were determined according to NIH guidelines as described in the Guide for the Care and Use of Laboratory Animals (8^th^ edition). Animals were anesthetized using isofluorane gas and the animals were euthanized through perfusion. Male Sprague-Dawley rats at 3 months of age were anesthetized, heparinized and perfusion fixed with 10% phosphate buffered formalin. Tissues were fixed at 4°C for 1 day and then stored in 20% ethanol in PBS thereafter until further processing. The tissues were then stained in 2.5% Lugol’s iodine solution for the following times based on the tissue types: sciatic nerve (2 days), heart (3 days) and brain (5 days). Prior to imaging the samples were rinsed in PBS and imaged semi-dry.

### X-ray Micro Computed Tomography

The tissues were imaged at the University of Texas High-Resolution X-ray Computed Tomography Facility (UTCT). For the sciatic nerve, the tissues were scanned on an Xradia MicroXCT 400 (Zeiss) using the 4X objective. The source-object distance was 37 mm, and the detector-object distance was 8 mm, resulting in 5.54 µm resolution. The X-ray source was set to 70kV and 10W and no X-ray prefilter was employed. A total of 1261 views were acquired over 360 degrees of rotation, at 2s/view. During reconstruction a beam-hardening correction of 3 was applied, and the data were byte-scaled to a range of -50 to 2800.

For the heart, the tissue was scanned with a custom scanner (North Star Imaging, Inc.). The scan used the Fein Focus High Power source with 130 kV and 0.14 mA (no filter). X-rays were detected with a Perkin Elmer detector using 0.25 pF gain, 1 fps, 1×1 binning, no flip, and a source to object distance of 143.0 mm. The overall source to detector distance was 1316.717 mm. A continuous CT scan, with no frames averaged, 0 skip frames, 3600 projections, 5 gain calibrations were taken. A 5 mm calibration phantom was used. The data range was set to -40.0 to 755.0 and had a beam-hardening correction of 0.4 was used. A post-reconstruction ring correction was applied using the following parameters: oversample = 2, radial bin width = 21, sectors = 32, minimum arc length = 8, angular bin width = 9, angular screening factor = 1. This resulted in a voxel size of 11.8 µm leading to 1961 total slices on the transverse axis of the heart.

For the brain tissues, the tissue was scanned with a custom scanner (NSI). The scan used the Fein Focus High Power source set to 110 kV and 0.18 mA (no filter). X-rays were detected with a Perkin Elmer detector with 0.25 pF gain, 1 fps, 1×1 binning, no flip and a source to object distance of 137.5 mm. The overall source to detector distance was 1316.635 mm. A continuous CT scan with 2 frames averaged, 0 skip frames, 3600 projections and 5 gain calibrations were taken. A calibration phantom of 0.762 mm was used. The data range was set from -40.0 to 695.0 and had a beam-hardening correction of 0.3. A post-reconstruction ring correction was applied using the following parameters: oversample = 2, radial bin width = 21, sectors = 64, minimum arc length = 2, angular bin width = 9, angular screening factor = 2. The resulting voxel size was 10.5 µm leading to 1989 slices on the coronal axis of the brain.

### Convolutional Neural Network and Visualization

Using the Deep Learning Tool in the Dragonfly 4.1 software (Object Research Systems, Inc.), we developed a binary segmentation model to segment the blood vessels of the three tissues following a standard U-Net architecture. The convolutional neural network model parameters are outlined in **Table 1**. Using five max pooling layers and five upsampling layers, we were able to develop robust models of binary segmentation in the three tissues. The max pooling layers contained two subsequent 2D convolutions followed by the 2×2 max pooling operation. The upsampling layers contain an up-convolution, a concatenation with a skip connection and two subsequent 2D convolutions. Training data sets were taken randomly throughout the CT slices. The selected slices were then manually segmented to form the training data set. As with previous iterations of the U-net convolutional neural network data augmentation was used to increase the robustness of the segmentation (**Table 1**). After training, the model was then used to segment the respective data set. This segmented data was then exported as a binary ROI and imported into the Avizo 9 software (ThermoFisher) for visualization. In Avizo 9, volume renderings and cutaway animations were created.

**Table 1.**
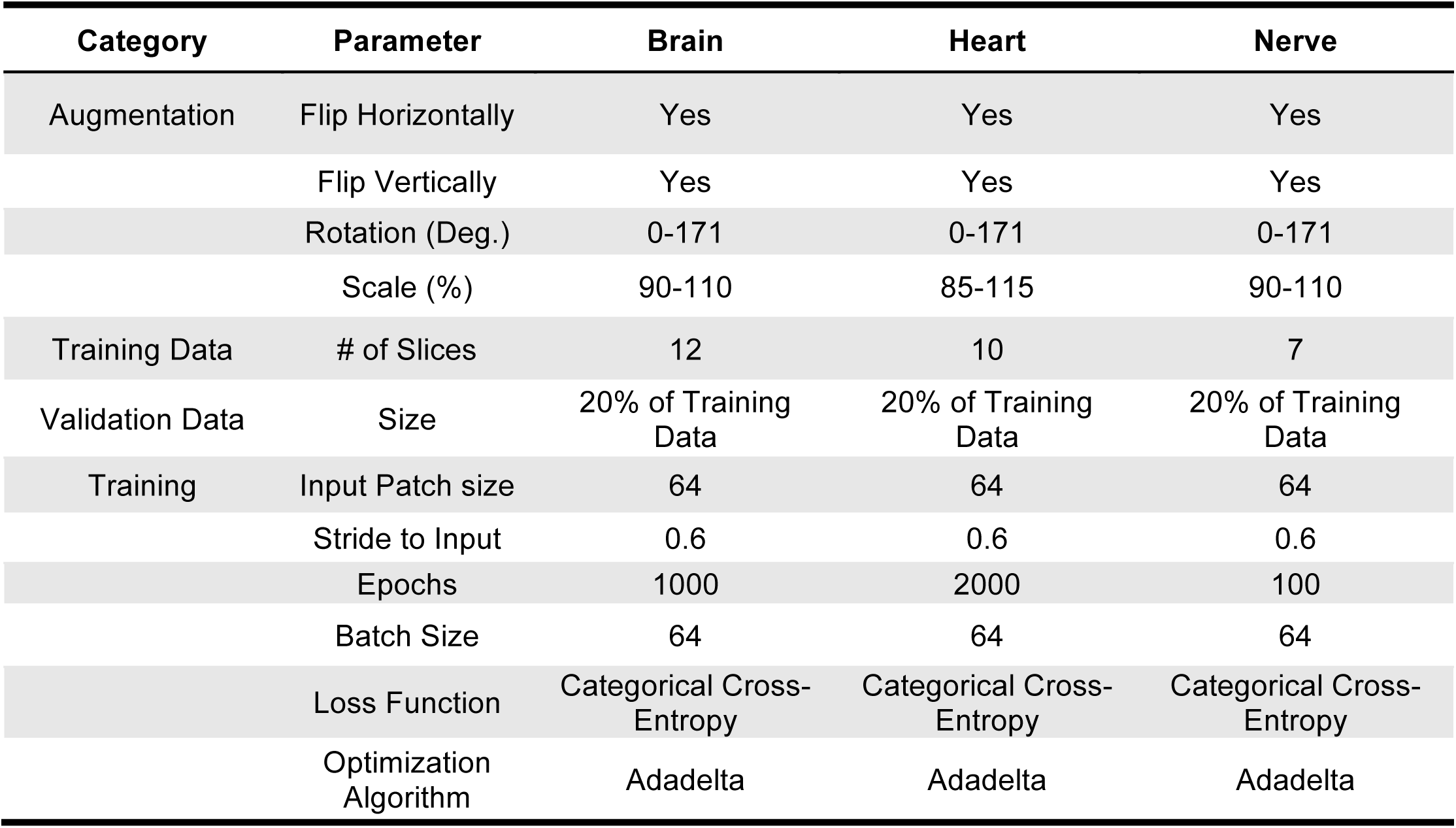
U-net Convolutional Neural Network Parameters.

### Cryosectioning and immunostaining

Following imaging, the samples were washed in 20% ethanol in PBS with changes every 12 hours for 3-5 days to remove the iodine staining. Tissues were then cryoprotected with treatment with progressive sucrose solutions increasing from 0% to 30% sucrose in PBS before being frozen in isopentane chilled with liquid nitrogen. A tissue matrix and cryotome blade was then used to cut subsections of the tissue. The samples were then sectioned using a cryotome keeping accurate track of the tissue depth to register with the Micro CT imaging. The sections were then dried at 60°C for 30 minutes before being stored at -20°C until immunostaining. The cryosectioned slides were then warmed to room temperature for 20 minutes before being rehydrated in PBS for 5 minutes. Antigen retrieval was performed using citric acid buffer (pH 6.0) was performed for 20 minutes in a 95°C water bath. After the slides were cooled the sections were permeabilized with 1% FBS and 0.4% Triton x-100 in PBS for 5 minutes. After washing with PBS, the slides were blocked in PBS with 20% FBS at room temperature for 45 minutes. The slides were then incubated with the primary antibody overnight at 4**°**C. The primary antibodies used were PECAM-1 (1:100 dilution, Abcam) and either beta III tubulin (1:100 dilution, Abcam) for the sciatic nerve sections, Troponin T (1:100 dilution, Abcam) for heart sections, and NeuN (1:100 dilution, Abcam) for brain tissues. The next day the slides were washed three times with 1% BSA in PBS. Then the slides were incubated with Alexafluor 488 and 594 (Abcam) dye conjugated secondary antibodies in 1% BSA in BSA at 1:500 dilutions. After a final wash, the slides were mounted with antifade mounting media and stored at 4**°**C until imaging. Images were taken with a Fluoview FV1000 Confocal Laser Scanning Microscope (Olympus).

## Results

### Micro CT visualization of sciatic nerve vasculature reveals blood vessel morphologies similar to measured confocal images

The overall flow for the staining and imaging procedure is shown in **Fig. 1**. Briefly, perfusion fixed samples were treated with an iodine solution that allows contrast between the vessels and surrounding tissues. The samples were then imaged using micro CT and the vasculature segmented using a convolutional neural network that was optimized for each tissue technique. We first tested this technique on sciatic nerve due the small sample size and relatively simple vasculature network structure in comparison to the other tissue types. After the nerve CT images were reconstructed, the CT image stack was cropped to a relatively small pixel volume (317 × 317 × 955 pixels) with a voxel size of 5.54 µm^3^ (**Fig. 2A**). We manually segmented the blood vessels on seven CT slices for our training data and subsequently segmented the rest of the nerve (**Fig. 2B**). After processing, the CT slices were reconstructed via a maximum intensity projection, allowing us to visualize the 3D morphology of the soft nerve tissue (**Fig. 2C**). We then created a volume rendering of the segmented blood vessels (**Fig. 2D**). After soaking the tissue in several changes of PBS for 3 days to remove the Lugol’s solution, the nerve was sectioned with a tissue matrix to locate a specific region identified in (**Fig. 2E.**), cryosectioned, and stained for beta III tubulin and PECAM-1 (**Fig. 2F**). Here we show a registration of the blood vessels segmented via our convolutional neural network and the histological result. The Lugol’s solution was shown to have no deleterious effects on the staining of nervous tissue. We measured blood vessel radii of both the CT images and confocal images and found a similar distribution of blood vessel size (**Fig. 2G**). Upon analysis of the radii of all the segments of blood vessels segmented by our convolutional neural network we show a distribution of mean blood vessel radii from the voxel limit of 2.77 µm to 13.47 µm (**Fig. 2H**).

**Figure 1.**
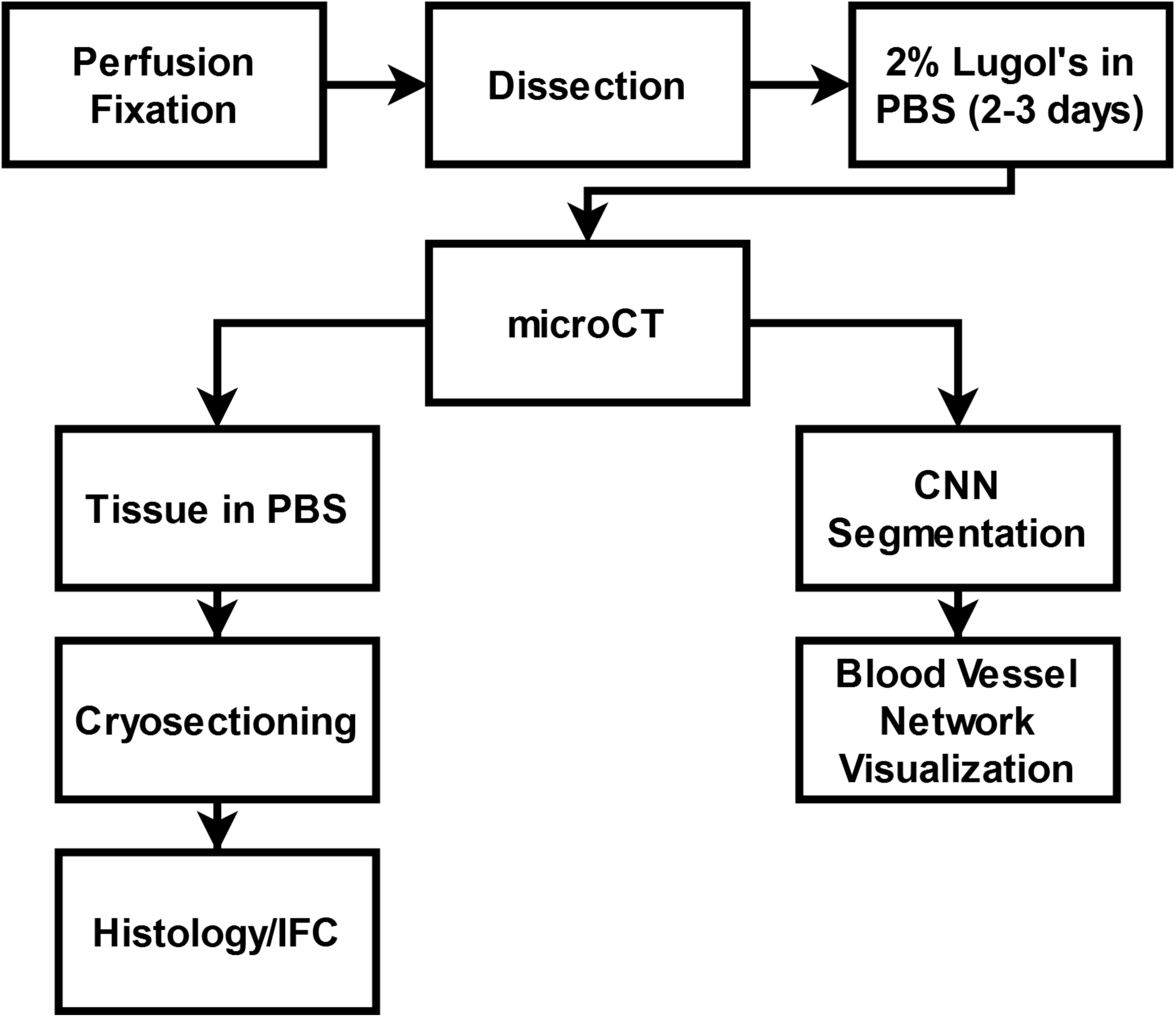
Methodology Overview. Following perfusion fixation and staining in Lugol’s solution, the tissue was scanned with microCT. The scans are then reconstructed and segmented with the convolutional neural network. Following imaging, the tissue was cleared of the contrast agent and then sectioned for histological preparations.

**Figure 2.**
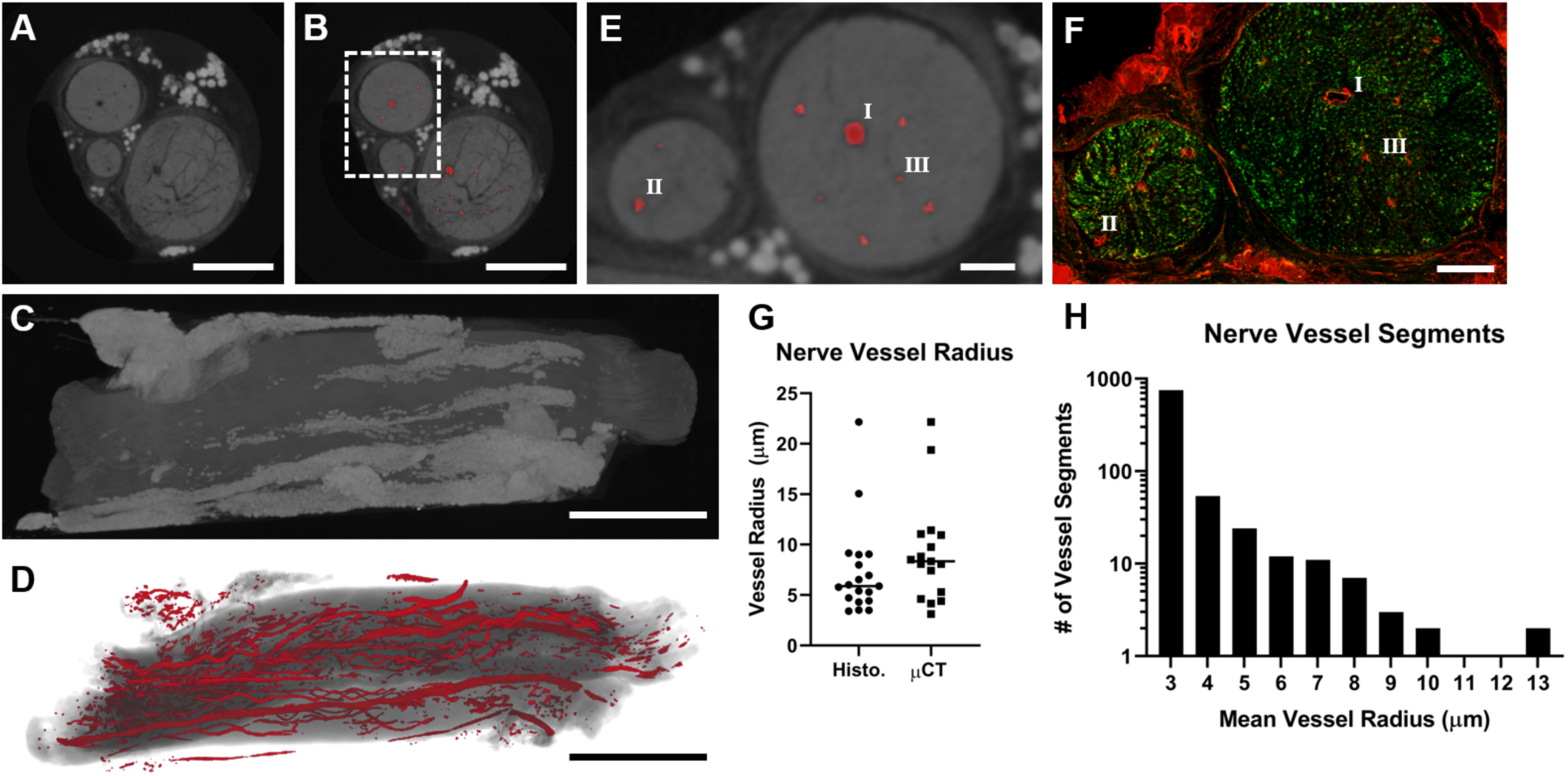
Application of the staining and segmentation applied to rat sciatic nerves. (A) The reconstructed micro CT images were cropped to a reduced volume. Scale bar = 500 µm. (B) The nerve was then segmented using our convolutional neural network. Scale bar = 500 µm. (C) The maximum intensity projection of the unsegmented micro CT stack. Scale bar = 1 mm (D) The volume rendering of the segmented blood vessels. Scale bar = 1 mm. (E) A close-up of the inset in B showing the segmented blood vessels in greater detail. Scale bar = 100 µm. (F) The nerve was then subsequently sliced and stained to confirm the validity of the blood vessel segmentation. The roman numerals in E and F show the same areas of the sample. Scale bar = 100 µm. (G) A measurement of vessel radii on confocal and micro CT images revealed no significant difference in vessel radius. (H) The frequency of micro CT segmented blood vessels radii shows our technique can segment both vessels at the voxel size limit as well as larger blood vessels.

### Visualization of heart vasculature identifies larger vessels but cannot resolve microvasculature due to voxel limitations

The visualization of coronary vasculature is sessential to understand disease states in the heart and the use of micro CT imaging has provided researchers a powerful tool to explore these illnesses^15,16^. After tissue preparation and scanning, the micro CT image stack was cropped to a reduced volume (1208 × 1246 × 1961 pixels; **Fig. 3A**). The blood vessels were segmented with a convolutional neural network trained on 10 slices taken randomly throughout the CT image stack. The convolutional neural network was then tasked with segmenting the rest of the heart (**Fig. 3B**). To visualize the full network in three dimensions, created a maximum intensity projection for the heart (**Fig. 3C**) and superimposed a volume rendering of the segmented vasculature (**Fig. 3D**). To assess the accuracy of the vascular segmentation, we identified a small region in the left ventricle (**Fig. 3E**) and performed cryosectioning to the heart tissue in this area for troponin T and PECAM-1 (**Fig. 3F, G**). For larger vessels, the method was able to register and segment the vessels with a high degree of accuracy but some of the smaller vessels were not captured by the method (**Supplemental Fig. 1** and **Supplemental Fig. 2**). The primary limitations for these small vessels came from the voxel size, which was dependent on the field of view needed to image the organ, and the contrast of the vasculature relative to the surrounding tissue.

**Figure 3.**
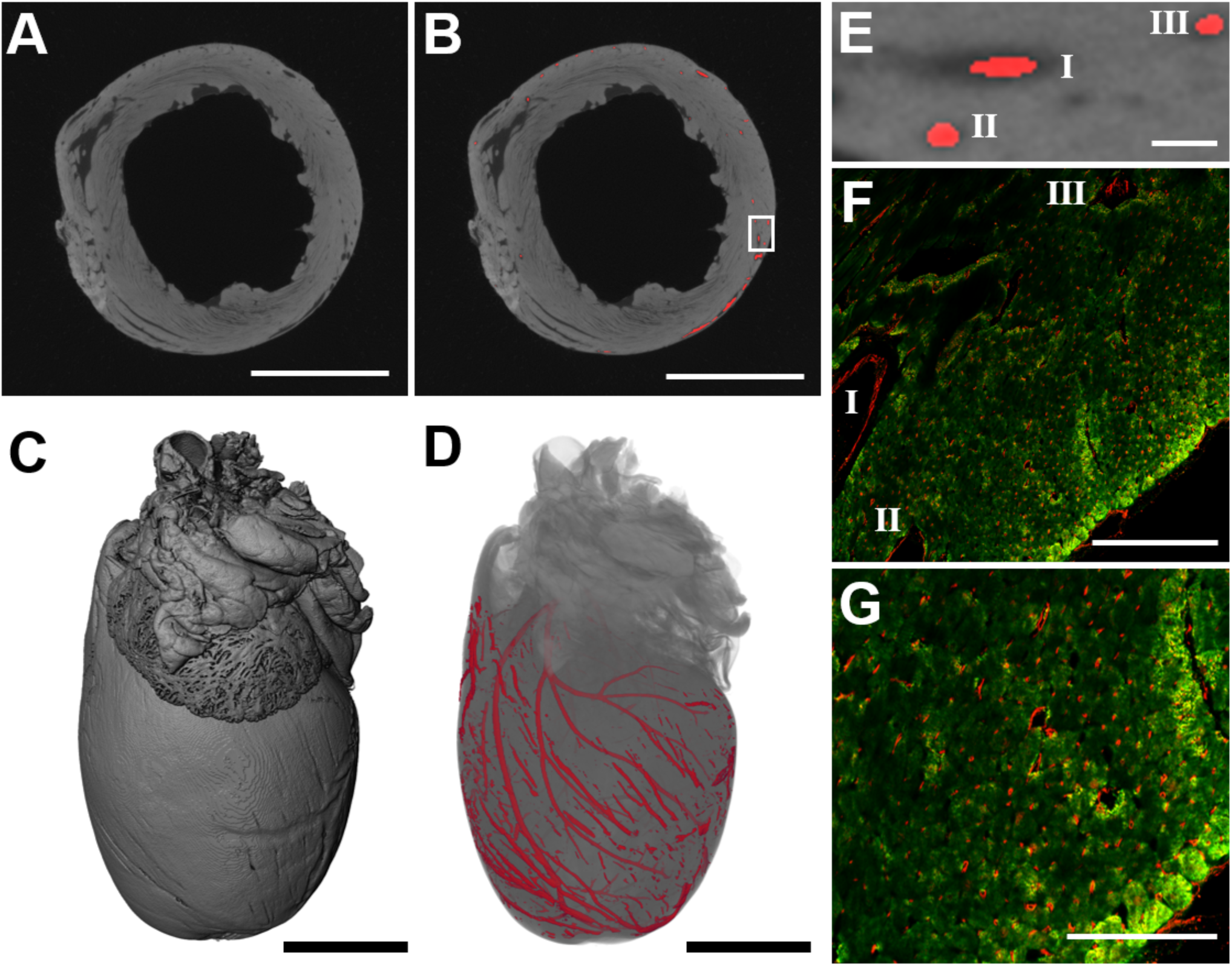
Application of the staining and segmentation method applied to a rat heart. (A) The reconstructed micro CT images were cropped to a reduced volume. Scale bar = 5 mm. (B) The heart was then segmented using the convolutional neural network. Scale bar = 5 mm. (C) The maximum intensity projection of the unsegmented micro CT stack. Scale bar = 2.5 mm. (D) The volume rendering of the segmented blood vessels. Scale bar = 2.5 mm. (E) A close-up of the inset shown in part (B). Scale bar = 200 µm. (F) The heart was then subsequently sliced and stained to confirm the validity of the blood vessel segmentation. The roman numerals in E and F show the same areas of the sample. Scale bar = 200 µm. (G) A magnified view the confocal image in (F) showing the presence of microvasculature that was not segmented by the convolutional neural network due to the size falling under the voxel size. Scale bar = 100 µm.

### The visualization of brain vasculature provides a general picture of the cerebral vessel network

To test the limitations of the method we also examine the ability of the technique to characterize the complex vascular network of the brain. The micro CT image stack was cropped to a reduced volume (1431 × 969 × 1989; **Fig. 4A**). A convolutional neural network was trained on 12 slices and again was tasked with segmenting the rest of the micro CT stacks (**Fig. 4B**). A maximum intensity projection was created for the brain (**Fig. 4C**) and a volume rendering of the segmented vasculature was superimposed onto this structure (**Fig. 4D**). The Circle of Willis and other vascular landmarks were visualized by the method. As with the heart, the larger voxel size needed to image the organ led to a loss of identification of some of the small microvessels. However, the technique captures a substantial number of descending vessels, including 10,465 individual segments (**Supplemental Figure 3**). The segmentation technique captured a wide range of mean blood vessel radii from the resolution limit up to 62.7 µm. We sectioned a portion of the cortex (**Fig. 4E**) and stained for NeuN and PECAM-1, demonstrating the maintenance of tissue epitopes in the brain after contrast-enhanced micro CT processing.

**Figure 4.**
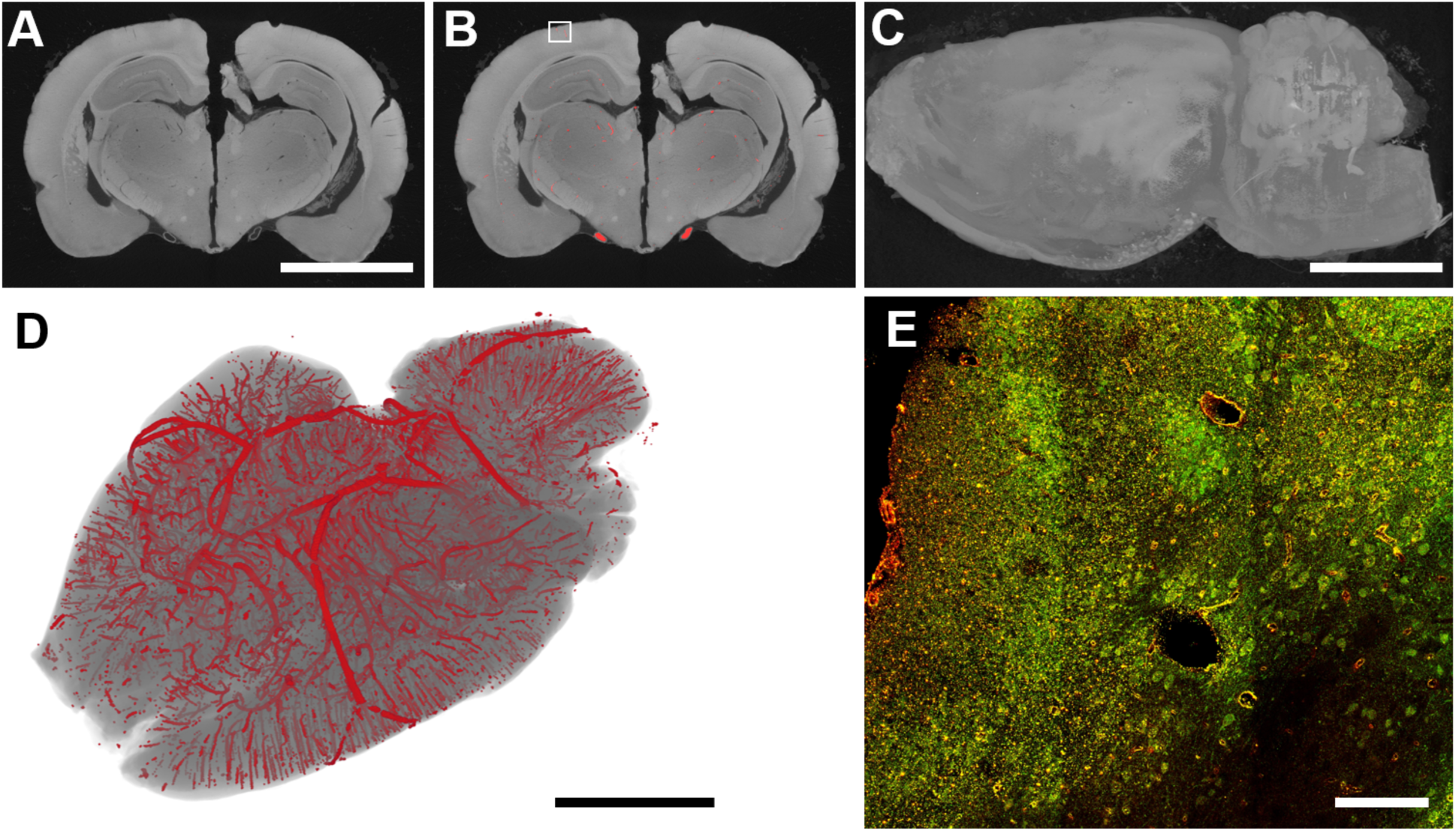
Application of the staining and segmentation method applied to a rat brain. (A) The reconstructed micro CT images were cropped a reduced volume. Scale bar = 5 mm. (B) The nerve was then segmented using our convolutional neural network. (C) The maximum intensity projection of the unsegmented micro CT stack. Scale bar = 5 mm. (D) The volume rendering of the segmented blood vessels. Scale bar = 5 mm. (E) The nerve was then subsequently sliced and stained to demonstrate the maintenance of epitopes through the method. Scale bar = 100 µm.

## Discussion

Convolutional neural networks have emerged as powerful segmentation tools for biomedical images where researchers may want to rapidly scan and identify regions of interest in huge data sets^12-14^. U-net convolutional neural network architectures require a much smaller set of training images, which was traditionally a bottleneck for biomedical image segmentation^17^. Furthermore, U-net based segmentation tools are widely available and can be employed in many different software environments, including Dragonfly^18^ and ImageJ^19^, making them widely accessible. Here, we demonstrate the application of a convolutional neural network for segmenting vascular networks in Micro CT images obtained using a reversible contrast staining procedure^20-22^. With this technique, tissues can perfusion fixed and processed with standard methods compatible with subsequent histological analyses. The method provides insights into vasculature in the whole tissue and can serve augment traditional analyses of vascular network performed using histology. The major limiters of the method used here included the voxel size obtainable by the micro CT imager for the desired field of view and the tissue specific contrast obtainable with the reversible iodine staining technique. For a small tissue sample imaged with small voxel size and high tissue to vessel contrast, as in the case of the sciatic nerve, there was a high degree of correspondence with the registered histology sections. With larger voxel sizes and less contrast with the surround tissues, as in the case of the heart and brain, the method works well with only for the larger vessels with the vascular network. Despite these limitations, the ability to visualize the overall vascular tree in three dimensions provides important information that can be used to better understand vascular network remodeling in the entire tissue and guide targeted analyses of the tissues by histology. In addition, the overall simplicity of the technique, nondestructive nature and relatively short processing time make the method easy to incorporate into the experimental pipelines of studies on vascular network remodeling.

## Acknowledgements

The authors gratefully acknowledge funding through the American Heart Association (17IRG33410888), the DOD CDMRP (W81XWH-16-1-0580; W81XWH-16-1-0582) and the National Institutes of Health (1R21EB023551-01; 1R21EB024147-01A1; 1R01HL141761-01) to ABB.

## Disclosures

None.

## Supplemental Figure Legends

**Supplemental Figure 1.**
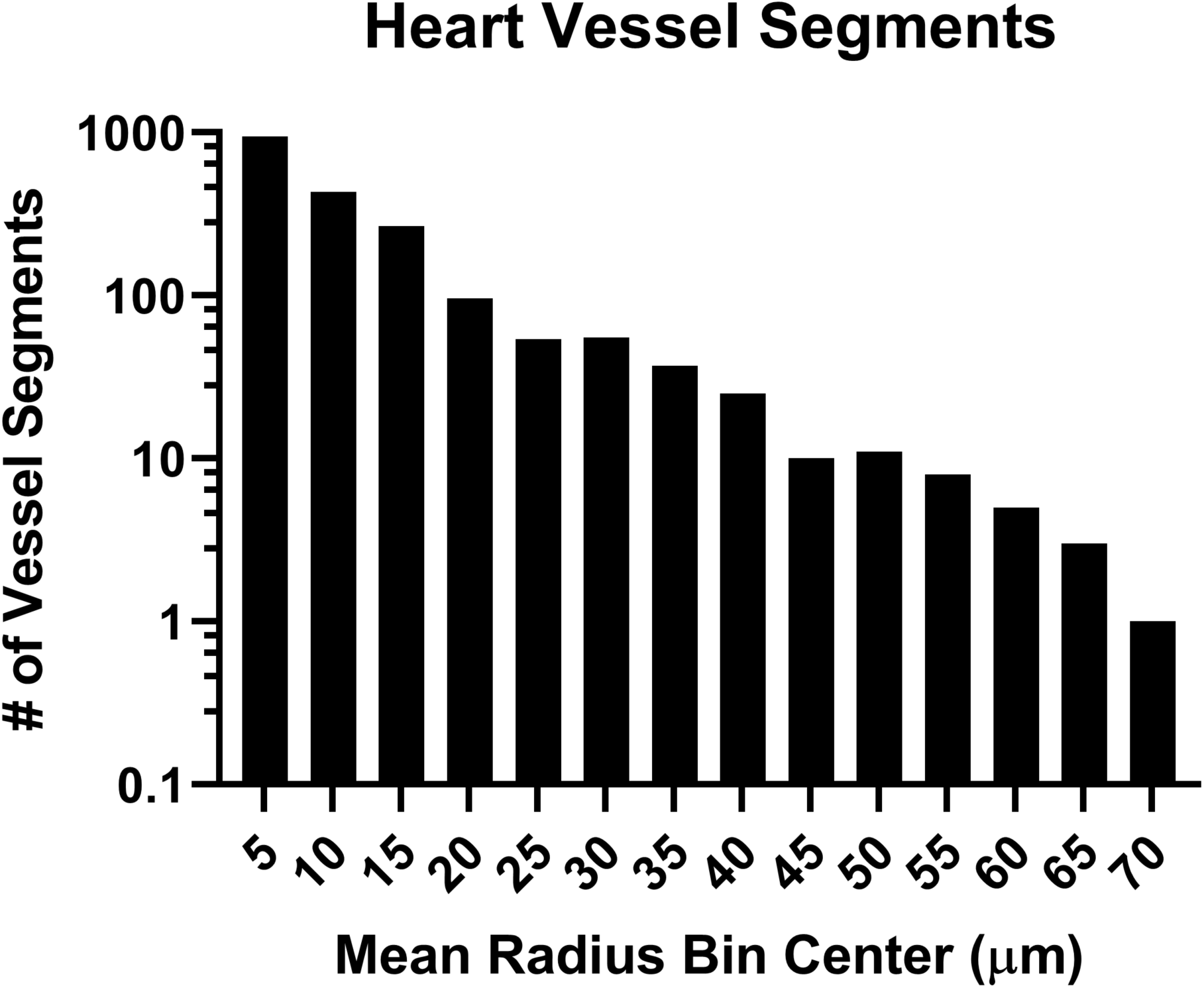
A histogram of the blood vessel radii segmented from the heart showing a wide distribution of mean vessel sizes from the voxel limit of 5.9 µm to 72.9 µm.

**Supplemental Figure 2.**
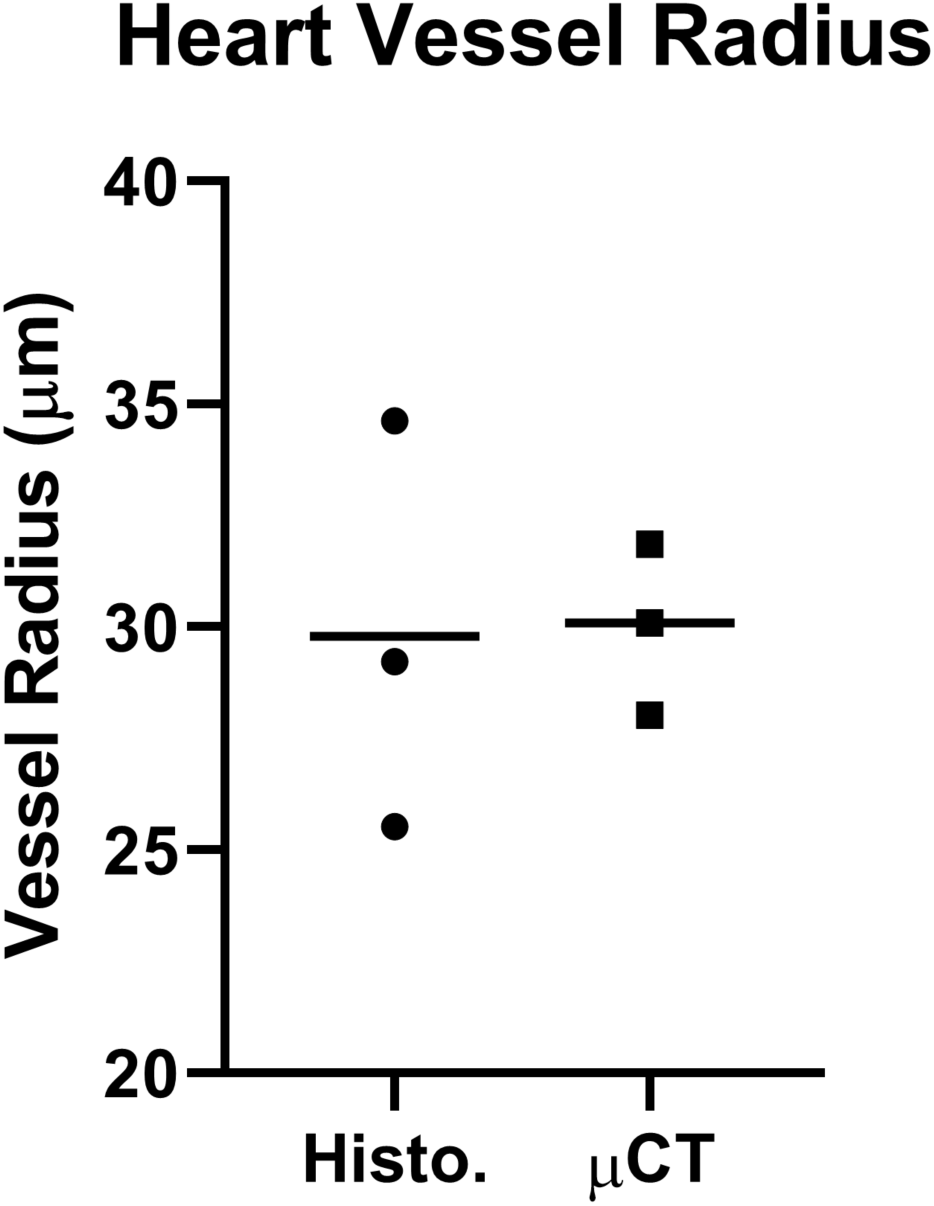
A comparison of the measurements of the three vessels indicated in Figure 3.

**Supplemental Figure 3.**
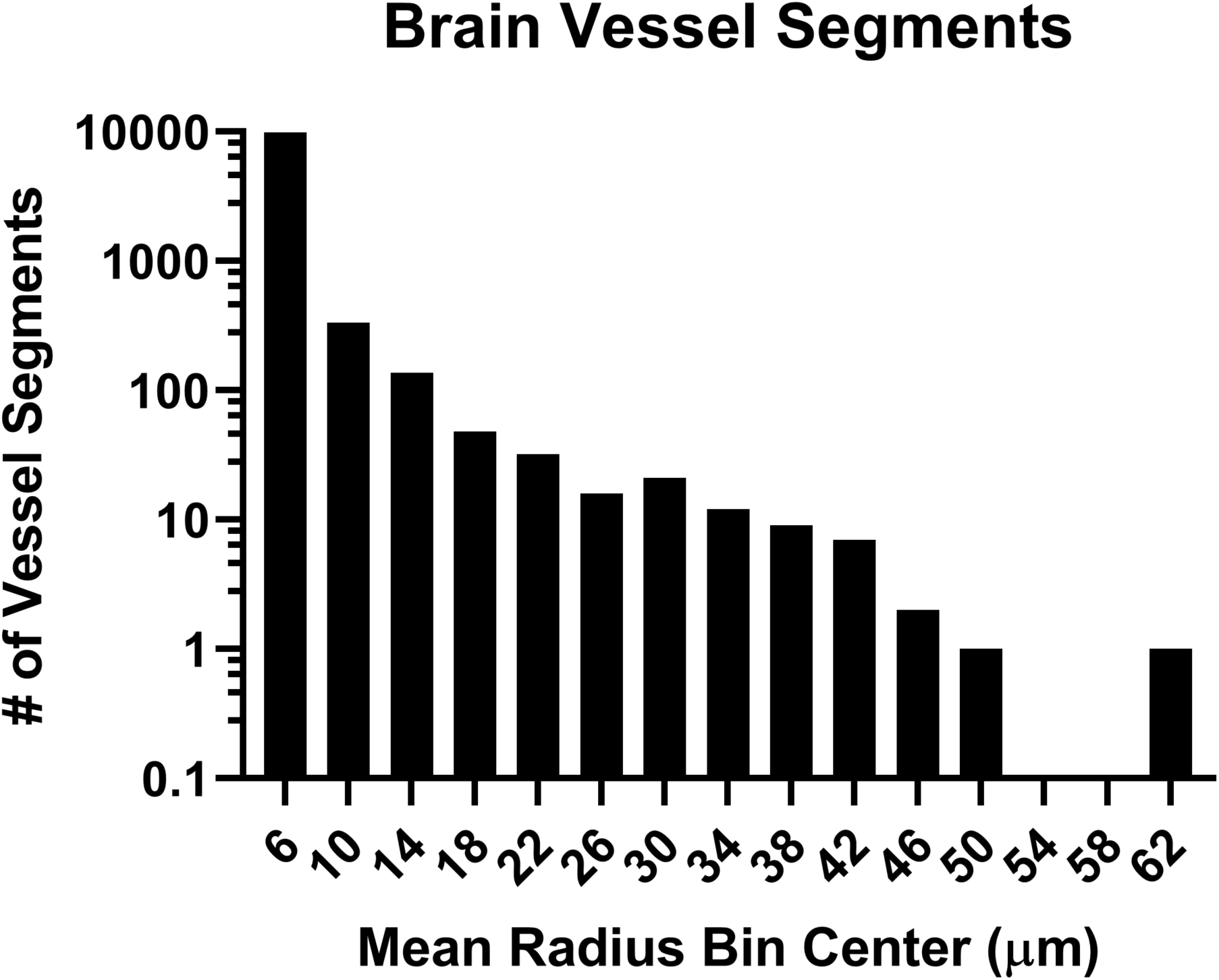
A histogram of the blood vessel radii segmented from the brain. The segmentation technique captured a wide range of mean blood vessel radii from the resolution limit up to 62.7 µm.

